# Fixed-target time-resolved crystallography at XFELs: the scourge of light contamination but reduced sample consumption

**DOI:** 10.1101/2023.12.12.571218

**Authors:** Guillaume Gotthard, Andrea Flores-Ibarra, Melissa Carrillo, Michal W Kepa, Thomas J Mason, Dennis P. Stegmann, Bence Olasz, Magdalena Pachota, Florian Dworkowski, Dmitry Ozerov, Bill F. Pedrini, Celestino Padeste, John H Beale, Przemyslaw Nogly

## Abstract

X-ray free electron laser (XFEL) light sources have allowed for the rapid growth of time-resolved structural experiments, which provide crucial information on the function of biological machines and their mechanisms. We set out to commission the SwissMX fixed-target sample delivery system at the SwissFEL Cristallina experimental station using the PSI developed MISP-chip for pump-probe time-resolved experiments. To characterise the system, we used the light-sensitive protein crystals of the Light-Oxygen-Voltage domain 1 (LOV1) from *Chlamydomonas reinhardtii*. Using different experimental settings, the adjacent-well light contamination was carefully assessed, indicating that it is crucial to control the light scattering from solid supports otherwise significant contamination can occur. However, our results show that, after the initial experiments and parameter refinement, the opaque MISP-chips are suitable for pump-probing a light-sensitive protein. This crystallographic experiment also probed the sub-millisecond structural dynamics of the LOV1 and indicated that at Δt=10 μs the covalent thioether bond is already established between the reactive Cys57 and FMN cofactor. This experiment validated the crystals to be suitable for in-depth follow up studies of the still poorly understood signal transduction mechanism. Importantly, the fixed-target delivery system also permitted a tenfold reduction in protein sample consumption compared to the most successful system used at XFEL, the high-viscosity extruder. This development creates the prospect of an exciting increase in XFEL project throughput for the field.

## Introduction

The emergence of serial crystallography has greatly benefited time-resolved macromolecular crystallography (MX). Developments in serial femtosecond crystallography (SFX) and serial synchrotron crystallography (SSX) enable even more experimental routines in the field and, altogether, have resulted in an expansion of time-resolved science. Although new triggering methods such as substrate mixing are continuously evolving, the optical laser remains the pump of choice for many pump-probe experiments. This is unsurprising given the time-resolved community’s early and ongoing focus on light-sensitive proteins. However, optical lasers can also be used to release photocages (Schulz *et al*., 2022), activate biological switches (Wranik *et al*., 2023), and induce temperature jumps in non-photoactive targets (Wolff *et al*., 2023) beyond the scope of light-sensitive proteins.

The very high peak brightness in X-ray free-electron laser (XFEL) sources that spurred the “diffraction before destruction” methodology (Wilmanns, 2000, Neutze *et al*., 2000, Chapman *et al*., 2011), creates the need for constant sample replenishment and high-performance sample delivery systems (also beneficial for synchrotron sources). The most widely used delivery systems include gas-focusing nozzles (DePonte *et al*., 2009), electrospun liquid microjets (Sierra *et al*., 2012) and the high-viscosity extruders, which work for both membrane and soluble proteins (Weierstall *et al*., 2014, Fromme *et al*., 2015, Nogly *et al*., 2015, Botha *et al*., 2015). Together these technologies have resulted in many novel insights into protein dynamics with the use of time-resolved SFX (TR-SFX). However, the sample efficiency of all jet-based methods is reduced when performing pump-probe measurements. During photoexcitation in a jet, the distance within the jet between adjacent XFEL shots needs to be safely larger than the width of the pump pulse (Smolentsev *et al*., 2013) to preclude shot-to-shot light contamination. The increased jet speed means higher sample consumption. Preparations for experiments often require months of work invested in purifying and crystallizing sufficient protein quantities. Therefore, any improvement in sample efficiency is of great interest to the structural biology field.

An alternative to jet-based systems is a fixed-target delivery system where protein micro-crystals are immobilized on a solid support or chip (Zarrine-Afsar *et al*., 2012). These techniques emerged later than their jet-based cousins but have proved to be robust and a user-friendly sample delivery system. Fixed-targets were first applied to SFX at LCLS where a micro-crystal slurry was applied to a silicon nitride membrane and rastered through in step with the XFEL pulse (Hunter *et al*., 2014). Since then, a large variety of different supports have emerged. These can be roughly grouped by whether they have apertures, and if these apertures need to be aligned to the beam (Carrillo *et al*., 2023). For these aperture-aligned fixed-targets, the key development was the coupling of precision silicon fabrication technique (Oghbaey *et al*., 2016) to precise stage motion and alignment strategies (Sherrell *et al*., 2015). The three prominent examples of these apertured targets are the HARE chip (Mehrabi *et al*., 2020) the Oxford chip (Ebrahim *et al*., 2019) and the Micro-Structured Polymer (MISP) chip (Carrillo *et al*., 2023). Examples of the raster-based targets are the NAM-based sample holder (Park *et al*., 2020, Nam *et al*., 2021), the Advanced Lightweight Encapsulator for Crystallography (ALEX) mesh holder (Sherrell *et al*., 2022), and the MPI sheet-on-sheet (SOS) chip (Doak *et al*., 2018). Solid support allows for a precise positional control that results in remarkable decreases in sample use without the need for gas flow or electrodes (Hunter *et al*., 2014, Mueller *et al*., 2015, Doak *et al*., 2018).

Given the utility of the fixed-targets and optical lasers for time-resolved crystallography, chip developers must ensure that optical lasers are compatible with their new tools. The significant issue in the case of fixed-targets is the precise illumination of a given crystal with an optical laser while leaving all the other crystals dark. This problem is not as trivial as it may first appear, and if not properly checked, it can lead to significant contamination of both the interleaved “light” and “dark” diffraction patterns, preventing the correct interpretation of relevant structural changes. In jet and extruder-based delivery systems, the oncoming crystals are safely housed in the dark enclosure of the jetting device and are delivered down a single axis, but this is not the case with fixed-target delivery [Figure 1(*a*)]. Fixed targets achieve time and sample efficiency by increasing the number of potential crystal locations on the chip face in a 2D array. The entire sample is loaded onto the target and is potentially reachable by the optical laser [Figure 1(*b*)]. Since the adjacent crystals are distributed in two dimensions, light contamination can propagate much further from the desired time delay [Figure 1(*c*)]; tens of milliseconds or even seconds depending on the location of contamination. This elongation of the probed domain will dilute the desired point of interest and probably create a large ensemble of different intermediates. Ultimately, making any subsequent analysis very challenging or, most likely, impossible.

**Figure 1.**
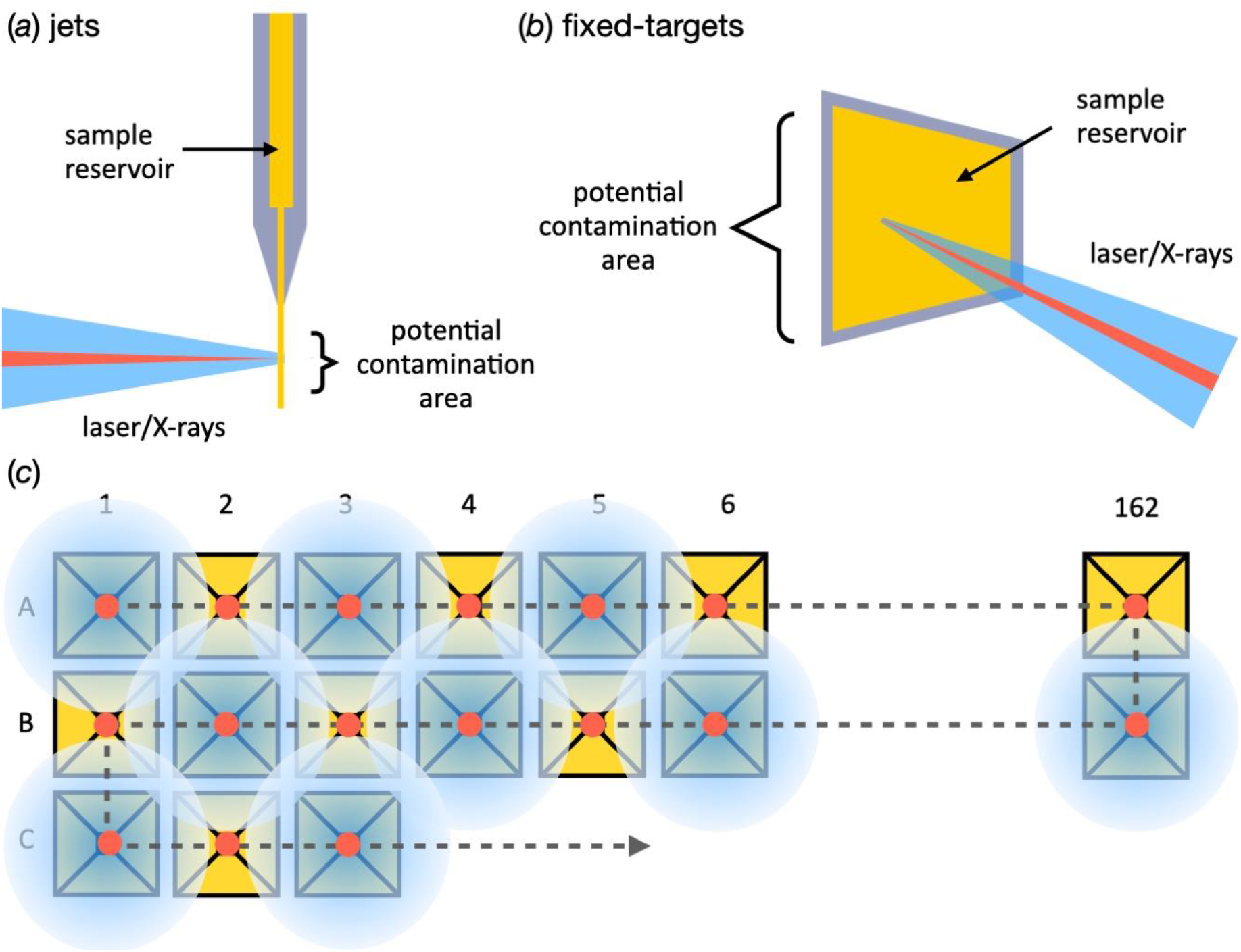
Practical differences between potential regions of light contamination in fixed-targets and jets. (*a*) and (*b*) A schematic of pump-probe in jets and fixed-targets, respectively. The sample reservoir in the fixed-targets is completely accessible to pump laser. (*c*) A schematic of a 50:50 interleaved light:dark setup in a MISP-chip. Importantly, light contamination in fixed-targets can happen both between consecutive apertures (10 ms at 100 Hz) and adjacent apertures, *i*.*e*., A1 to B1 or even C1. The contamination of these adjacent apertures give rise to much longer timescale contamination; A1 to B1 = 3.24 s. The sample area or cavities are depicted in yellow, the X-ray are shown in red, and the stage path is depicted as a dashed grey line. The pump laser is shown in blue.

XFELs are uniquely suited to the study of ultrafast dynamics on the femtosecond-microsecond scale; however, fixed-targets have yet to be exploited for time-resolved work at XFELs; although their utility has been proven for similar experiments at synchrotrons (Mehrabi *et al*., 2019). The serial crystallography with solid-support MX (SwissMX) endstation at the SwissFEL Cristallina experimental station (Paul Scherrer Institute, Villigen, Switzerland) and PSI developed MISP chips were designed to address this gap. However, extensive tests were obviously essential to validate these tools for optical laser-based pump-probe experiments. This work describes a commissioning experiment conducted in November 2022 at the SwissFEL Cristallina station, demonstrating the utility of the MISP chip (Carrillo *et al*., 2022) for TR-SFX. The aim was to find and understand the parameters that enable time-resolved pump-probe experiments using the SwissMX and MISP chips. To achieve this, a pump-probe 50:50 light:dark (hereafter referred to as interleaved-light and interleaved-dark, respectively) experimental routine with both transparent and opaque MISP chips was undertaken. The protein used in these experiments was the Light-Oxygen-Voltage (LOV) domain 1 from *Chlamydomonas reinhardtii* (*Cr*LOV1). *Cr*LOV1 is a light-sensitive protein domain that undergoes a long photocycle (∼200 s recycle time) and, therefore, is acutely sensitive to photoactivation from stray photons from adjacent cavities. A covalent thioether bond forms between the flavin mononucleotide (FMN) cofactor and Cys57 from the protein binding site after photoexcitation (Fedorov *et al*., 2003). These tests also served to benchmark the data collection making *Cr*LOV1 an ideal candidate with a distinct active state signature that can be used to assess adjacent-well light contamination.

## Material and methods

### Expression and purification

The expression and purification protocols have been described previously (Gotthard *et al*., 2023). Briefly, the expressed protein sequence was amino acids 16-133 of the Phot1 protein LOV1 domain from *Chlamydomonas reinhardtii*. The protein possess an N-terminal His-tag followed by a Factor Xa protease site. The expression was conducted in *Escherichia coli* BL21 DE3 using auto-inducible media (Studier, 2005). The expressed protein was purified by Ni^2+^ affinity followed by size exclusion chromatography. Fractions corresponding to the protein were pooled and concentrated to 10 mg·ml^-1^ for crystallization.

### Crystallization

Limited proteolysis with trypsin allowed a more reproducible and controlled crystallization process (Gotthard *et al*., 2023). Microcrystals were grown at 293 K in batch with a 2:1 protein to well condition ratio in 0.1 M Sodium cacodylate pH 6.5, 1.0 M Sodium citrate dibasic trihydrate. An average crystal size of 25 μm ∓ 7 μm was measured using a Leica microscope(Kaminski *et al*., 2022). The crystals concentration was measured using a cell counter and estimated to be 1.0-2.0 × 10^7^ crystals·ml^-1^.

### Chip mounting

Firstly, the crystal concentration was adjusted by diluting the micro-crystal slurry with crystallisation solution to a final concentration of 1-2×10^6^ crystals·ml^-1^. Subsequently, 400 μl of crystalline solution was pipetted onto a MISP-chip that was either made with transparent cyclic olefin polymer (COP) film or an opaque film made by mixing 10 % (w/w) carbon black with cyclic olefin copolymer (COC) (Carrillo *et al*., 2023). Once loaded, the MISP-chip was placed on a loading stage connected to a vacuum that served to remove the excess mother liquor and funnel the crystals into the wells. Filter paper was occasionally required to blot away any excess solution. After this, the chip was placed onto the MISP-chip holders (Carrillo *et al*., 2023) that sealed the chip inside two pieces of 6 μm Mylar® film and maintained the crystal hydration. They were then placed inside a darkened humidity chamber kept at 80 % relative humidity and transported to the beamline. X-ray data collection was performed at the Cristallina experimental station of the SwissFEL using the SwissMX endstation. Due to concerns over crystal hydration, only 5 chips were consecutively loaded and kept in the humidity chamber at a time. Chips were manually mounted from the humified chamber to the SwissMX. Collecting each set of 5 chips took approximately 50-60 minutes and all of this was done at room temperature (23-25 °C, depending on the location within SwissFEL).

### Beamline setup

Data were collected over a 24-hr period on November the 27^th^ 2022. The X-ray beam energy was 12.4 keV with a pulse energy at the sample position of approximately 50 μJ. The X-rays were focused using Kirkpatrick-Baez (KB) mirrors to a spot size of approximately 1.5 × 1.5 μm and the repetition rate was 100 Hz. The pulse width was approximately 35 fs root mean square (RMS). The diffraction data were recorded on a JUNGFRAU 8 Mpixel detector.

### Laser coupling

As of November 2022, the Cristallina station was not provided with a through-space connection to a pump laser. The SwissMX endstation was limited to a 70 m fiber connection to a nanosecond laser located in the SwissFEL laser room. Due to this constraint, the decision was taken to couple the laser to the sample position through the SwissMX on-axis-viewing (OAV) system. Such couplings are common on synchrotron protein crystallography beamlines and this solution offered the best compromise between the final achievable focus and the physical meshing with other instrumentation at the sample position.

For fixed-targets the key parameter in a pump-probe experiment is the laser focal-spot size. The efficiency of the fixed-target is dependent upon the number crystal locations that can be squeezed onto the chip-surface. For the apertured fixed-targets this means having a small pitch between adjacent cavities. The achievable laser focus, therefore, has a direct influence on the efficiency of the experiment as an increase in the diameter of the focus will necessitate an increase in the aperture pitch.

The laser spot size was estimated to be 50x50 μm^2^ based on its reflection from a piece of opaque COC film held at the OAV focus. The OAV was calibrated against known distances, but it is impossible to accurately infer from this the 1/e^2^ or full-width half-maximum (FWHM). The pulse energy was set to approximately 2 μJ.

### Pump-probe setup

X-ray data collection over the whole chip in a dark environment without any laser excitation was used as ‘reference’ for the calculation of the Fourier difference electron density maps (*F*_obs_^laser-off^). Light contamination from the transparent and opaque chips and SwissMX setup was assessed using two chip orientations: *open*, with the wider side of the cavity directed towards the laser, and *flat*, with the wider side of the cavity facing the detector. All pump-probe data were collected 50:50 interleaved light:dark, *i*.*e*., XFEL at 100 Hz, ns laser at 50 Hz, giving a laser pulse in every other aperture.

### Data processing

Serial data processing was performed using the CrystFEL version 0.10.1 software suite (White, 2019). Diffraction hits were identified using the peakfinder8 and XGANDALF (Gevorkov *et al*., 2019) algorithms with a hexagonal unit cell (a = b = 122, c = 46 Å). Peaks integration was performed using the three-rings methods in indexamajig with integration radii of 2, 3, and 5 pixels. Indexing rates were between 50 and 80 %. Interleaved-dark and the interleaved-light image lists were generated by the labelling images with a ‘laser-on’ event generated by the SwissFEL event master. This event is passed to the JUNGFRAU whilst data are collected and propagated with it thereafter. By following the laser events, the interleaved data could be indexed and integrated independently. Stream files were merged using partialator using the unity partiality model with a pushres option of 1.6-2.0 nm^-1^. *hkl* files were converted into *mtz* with *f2mtz* from the CCP4 suite(Winn *et al*., 2011). A resolution cut-off was applied when CC_1/2_ was falling below 30%. Dataset statistics are reported in Table 1.

**Table 1.**
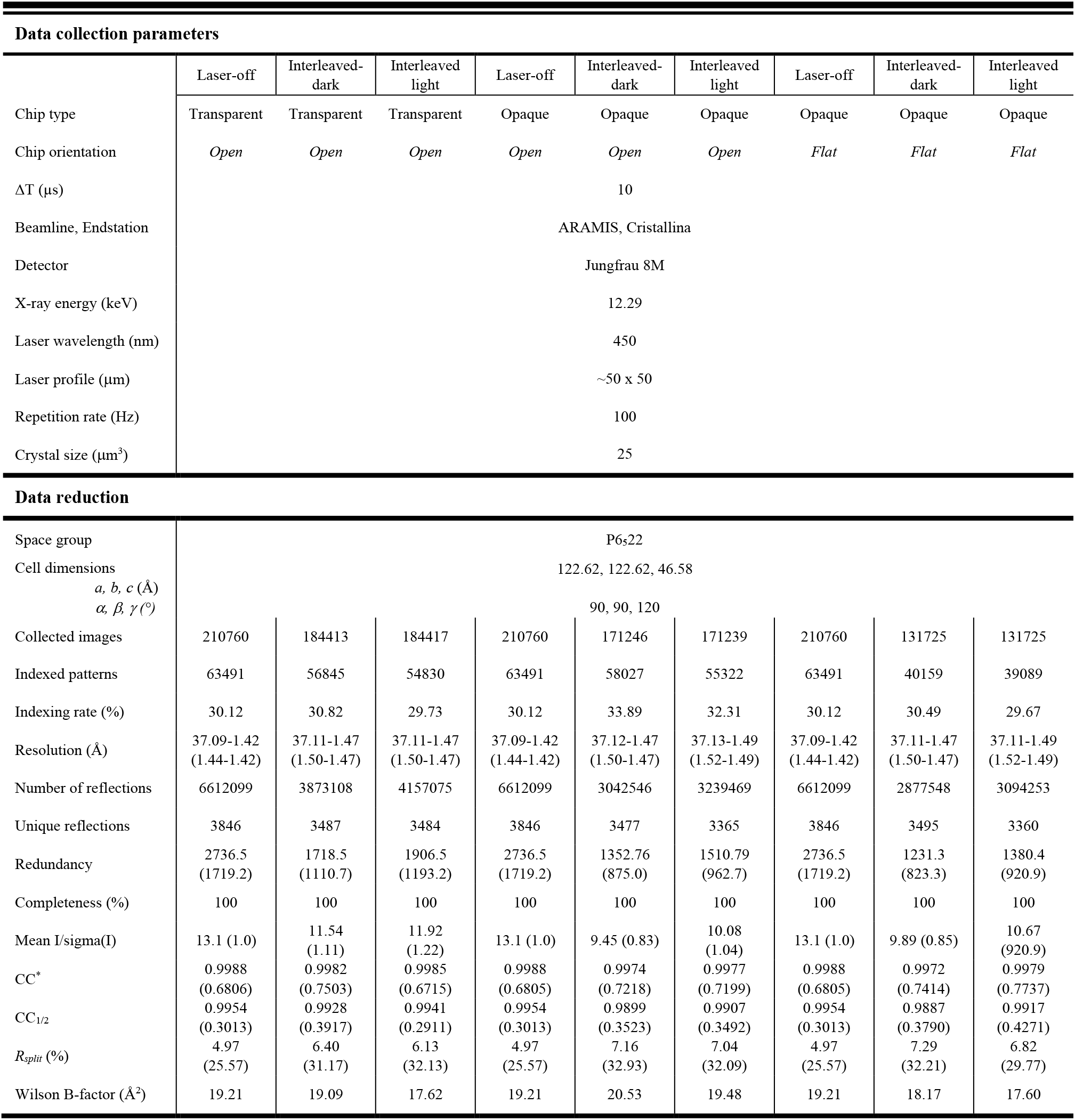
Data collection parameters and data reduction for the 1^st^ experiment.

### Isomorphous difference maps

Fourier difference electron density maps were calculated using the phenix.fobs_minus_fobs_map program from the Phenix suite (Liebschner *et al*., 2019). A resolution cut-off of 1.8 Å and a sigma cut-off of 3.0 were applied and the multiscale option was used to calculate the difference maps. The presence of contamination could be observed by subtracting data collected without laser illumination (*laser-off*) from the interleaved data (*interleaved dark or light*): *F*_obs_^interleaved-dark/light^ – *F*_obs_^laser-off^. Assuming no contamination can be observed, the signal from the *F*_obs_^interleaved-light^ – *F*_obs_^laser-off^ should also be the same as the interleaved difference map, *F*_obs_^interleaved-light^ – *F*_obs_^Fobsinterleaved-dark^. Figures were prepared using PyMOL (DeLano, 2020).

## Results and discussion

### Pump-probe with fixed-targets

Here, we present laser-triggered, pump-probe experiments using the SwissMX endstation and MISP-chips at the SwissFEL Cristallina experimental station. To translate samples, the SwissMX is equipped with two orthogonal linear stages (Parker) for *x* and *y* motions and two additional stages *z* and additional *x* motion (Standa). The MISP-chips have a reliable active area of 162 × 162 cavities totalling 26,244 apertures. All data from the chips were collected in a serpentine-like pattern. Figure 2 shows the final stage of the optical pump-laser coupling through the SwissMX OAV. The setup is current only used for in-air data collection. Scatter guards are, therefore, required to catch the beam from the OAV to the sample and from the sample to the detector face. The total exposed length to air is approximately 15 mm.

**Figure 2.**
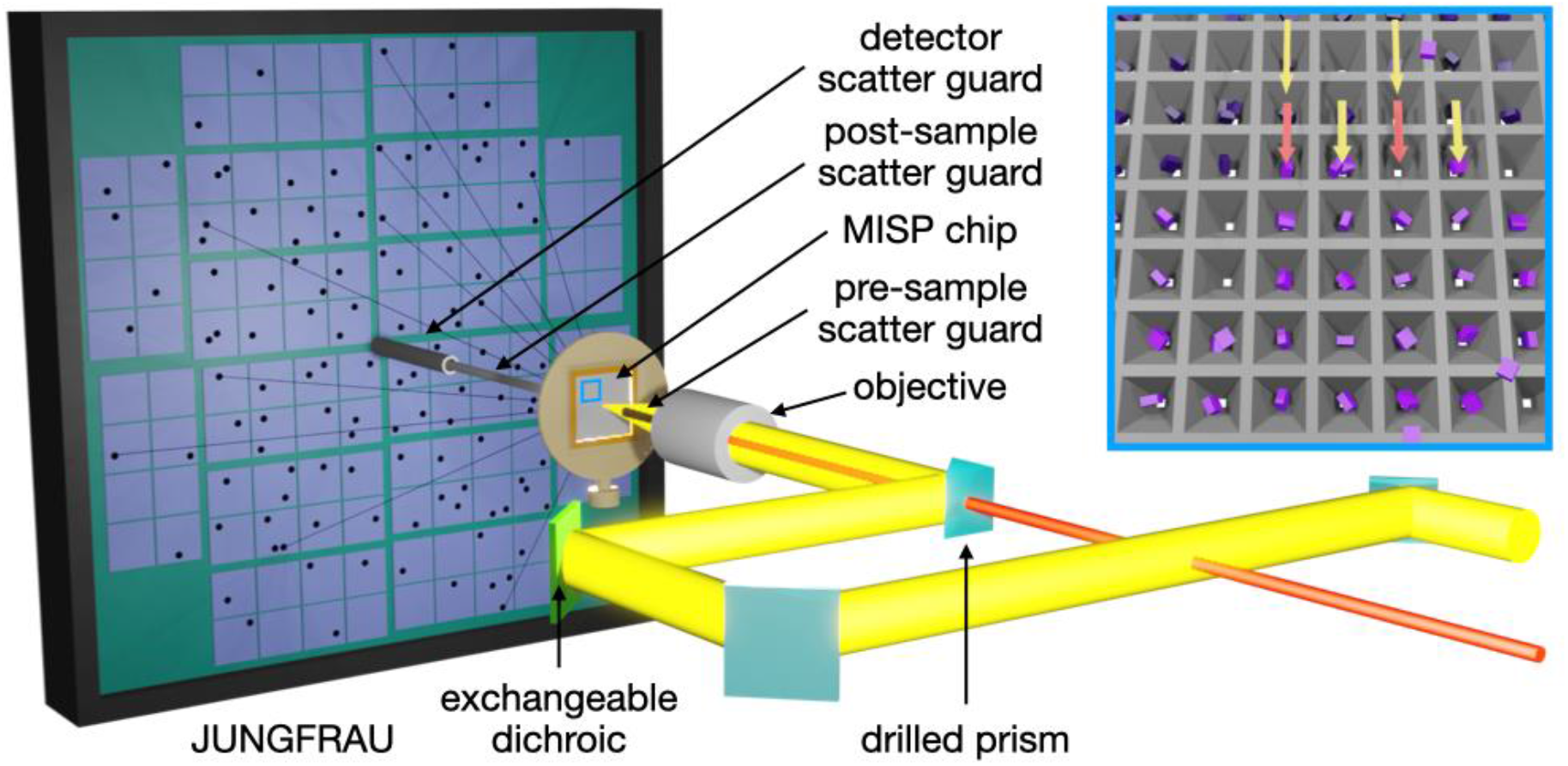
A schematic of the pump-probe experimental set-up at Cristallina, SwissFEL. The pump laser (yellow) is coupled to the sample position *via* the endstation OAV. An exchange dichroic mirror (light green) enables different pump wavelengths to be reflected whilst transmitting light for the chip alignment. The X-rays pass through the center of the drilled objective and prism of the final part of the laser coupling. Air scatter from the X-ray is minimized using pre-and post-sample scatter-guards. The blue box highlights an area of the chip showing a 50:50 interleaved light:dark scheme, where X-rays are delivered to every well and laser pump only to every other.

Data were collected on two different chip types, named *transparent* and *opaque* from their transparency to visible spectrum or lack thereof, and in two different orientations, *open* and *flat*. The transparent chips were made from commercially available 50 μm COP film, whereas the opaque chips were fabricated with an in-house cast film using COC pellets and the addition of carbon black powder (Carrillo *et al*., 2023)

Time-resolved spectroscopy experiments on *Cr*LOV1 used in this experiment indicated that it undergoes the formation of a covalent thioether bond between FMN cofactor and Cys57 at Δt=10 μs after photoexcitation (Kottke *et al*., 2003, Holzer *et al*., 2002), which then persist even late into the photocycle (Fedorov *et al*., 2003, Kottke *et al*., 2003) (Figure 3). Given the significant strength of expected signal in the Fourier difference maps for the covalent bond formation, this time delay was selected to test the suitability of the *Cr*LOV1 crystals for TR-SFX experiments and make use of their long photocycle, on the order of 200 s, as a light contamination indicator.

**Figure 3.**
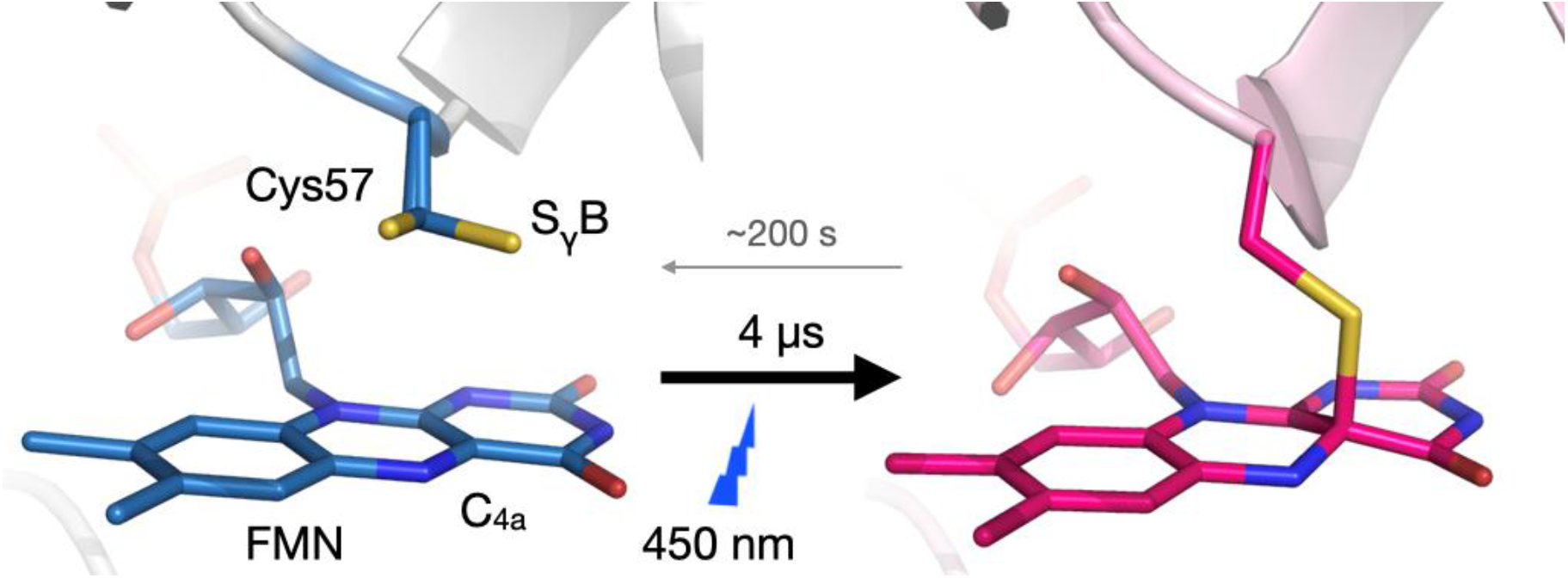
Thioadduct formation between *Cr*LOV1 and its FMN cofactor upon illumination. Crystal structures of *Chlamydomonas reindhartii* LOV1 domain in light and dark stationary states present two conformations of the binding-site cysteine (Cys57), where blue-light absorption readily causes after 10 μs the formation of a covalent bond between the flavin C(4a) and the thiol of Cys57. The reaction proceeds through an excited flavin singlet to a triplet state that then decays monotonically to the adduct in which this cysteine moves ∼1.5 Å closer to the FMN-C(4a) for adduct formation (Holzer *et al*., 2002, Kottke *et al*., 2003). Light absorption (and thus, thioadduct formation) then triggers rearrangements throughout the whole LOV domain. **(Gotthard *et al***., **2023)**

We employed a 50:50 interleaved light:dark experimental routine with the XFEL at 100 Hz and the pump-laser at 50 Hz. While a comprehensive analysis of the LOV-FMN light-activated structure is beyond the scope of this paper, it suffices to expect a photoactivated structure with the FMN-Cys covalent adduct formation in the laser-exposed wells. The unpumped wells should yield a dark state structure without the thioether bond signature. We also collected SFX data entirely without pump laser (*laser-off*) for reference as a “properly” dark state. Ultimately, the *laser-off* should be indistinguishable from the dark datasets in the interleaving TR-SFX experiment.

It is important to stress the importance of delivering light-contamination-free time-resolved data. Otherwise, the crystallographic data would represent multiple overlapping protein trajectories triggered by more than one pump laser pulse, impairing its correct interpretation. Importantly for the current work, the *Cr*LOV1 domain was specifically chosen for the fixed-target pump-probe commissioning since the 200 s resting state recovery time of the domain significantly exceeds the 10 ms interval between the consecutive XFEL pulses into the adjacent cavities of the chip. The 200 s recovery time also exceeds the time to image over half the chip, so potential adjacent column light contamination will also be observed.

In the quest to explore different experimental geometries of the fixed-target setup, two orientations of the chips with respect to the incidence pump laser and X-rays were tested. One with laser incidence coming from the wider side of the pyramidal cavity (*open* side) or coming from the narrower side (*flat* side). This option has been debated since the original use of apertured opaque fixed-targets for pump-probe experiments since this decision may have an impact on the excited fraction of the crystal (see Figure 4 left hand panel).

**Figure 4.**
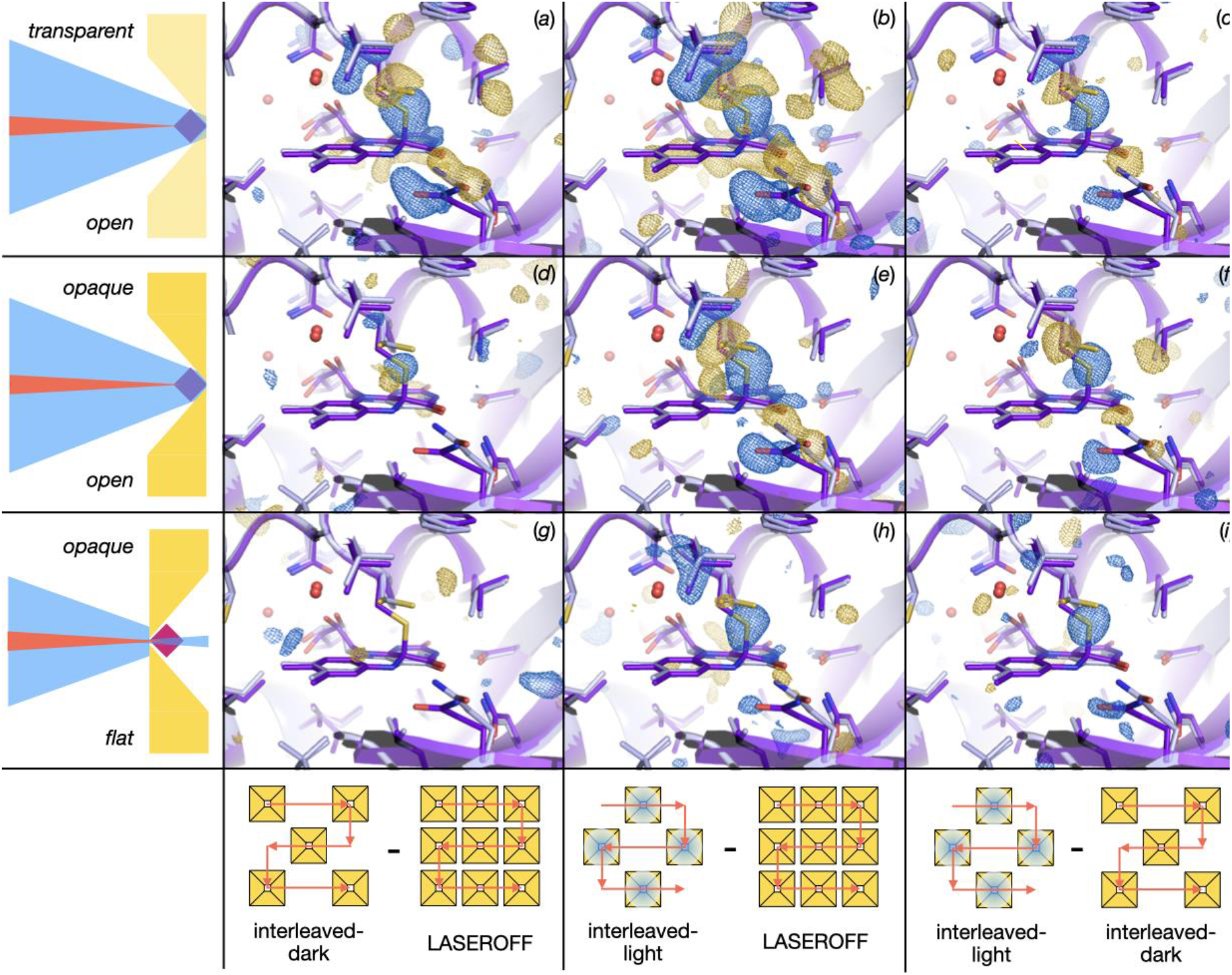
Chip placement for laser incidence and Fourier difference electron density maps for TR-SFX with *Cr*LOV1 at Δt=10 μs (first experiment). *Transparent chip with cavity facing the pump pulse* yields activation signal in all three maps corresponding to thioether bond formation between Cysteine and FMN: (*a*) *F*_obs_^interleaved-dark^ – *F*_obs_^laser-off^ (*b*) *F*_obs_^interleaved-light^ – *F*_obs_^laser-off^ (*c*) *F*_obs_^interleaved-light^ – *F*_obs_^interleaved-dark^ *Opaque chip with cavity facing the pump pulse* yields activation signal in all three maps: (*d*) *F*_obs_^interleaved-dark^ – *F*_obs_^laser-off^ (*e*) *F*_obs_^interleaved-light^ – *F*_obs_^laser-off^ (*f*) *F*_obs_^interleaved-light^ – *F*_obs_^interleaved-dark^ *Opaque chip with aperture facing the pump pulse* yields activation signal where expected while the lack of signal in the F_dark_ – F_laser-off_ indicates that in this setup light contamination was avoided: (*g*) *F*_obs_^interleaved-dark^ – *F*_obs_^laser-off^ (*h*) *F*_obs_^interleaved-light^ – *F*_obs_^laser-off^ and (*i*) *F*_obs_^interleaved-light^ – *F*_obs_^interleaved-dark^. Fourier difference electron density maps show positive density depicted in blue (atoms move in) and negative density depicted in gold (atoms move out) highlighting differences between datasets. The cartoon and sticks representation show in light grey the dark state of the protein with its FMN ligand adjacent to it but not covalently bound and in purple the photoactivated late photocycle intermediate for reference with FMN covalently binding to LOV. To the left side of the difference electron density maps there is a schematic representation of the chip (yellow) with a crystal sample (pink/blue) placement with respect to pump laser (blue) and XFEL pulses (red). The bottom panel shows the practical arrangement of the data from the wells contributing to the different maps.

### Assessment of contamination

The TR-SFX experiment was carried out with a Δt=10 μs between the laser pump and X-ray probe pulses. Fourier difference electron density maps were evaluated in terms of successful photoactivation and potential light contamination in the nearby wells. The transparent chip setup yielded positive signal indicating thioether bond formation in *F*_obs_^interleaved-dark^ – *F*_obs_^laser-off^ [Figure 4(*a*)], *F*_obs_^interleaved-light^ – *F*_obs_^laser-off^ [Figure 4(*b*)] and *F*_obs_^interleaved-light^ – *F*_obs_^interleaved-dark^ [Figure 4(*c*)]. While it presence of features characteristic of the light activated state, the light signal present in the *F*_obs_^interleaved-dark^ – *F*_obs_^laser-off^ indicates that each pump laser pulse did not exclusively photoactivate a single well of the chip but that the light reached the adjacent wells leading to light contamination. Although light contamination was more likely in the transparent chips, the level of its pervasiveness was not expected and was interesting to observe that the signal for the *F*_obs_^interleaved-dark^ – *F*_obs_^laser-off^ was as strong as for the *F*_obs_^interleaved-light^ – *F*_obs_^interleaved-dark^ maps. The light contamination was likely due to the transmission of the laser light orthogonally through the chip from the scattering source, either an interaction with the crystal or chip.

Next, we performed tests using the opaque MISP-chips in two orientations. The *open* orientation, which allows the most light to reach the sample and maximizes the excited fraction of molecules, again yielded positive signal indicating the thioether bond formation in the *F*_obs_^interleaved-dark^ – *F*_obs_^laser-off^ [Figure 4(*d*)]. Although yielding significantly weaker signals than in the case of transparent chips, contamination is still evident. A similar signal was also present in the *F*_obs_^interleaved-light^ – *F*_obs_^laser-off^ [Figure 4(*e*)] and *F*_obs_^interleaved-light^ – *F*_obs_^interleaved-dark^ [Figure 4(*f*)].

However, in the alternative chip orientation, with the *flat* side of the cavity now facing the pump laser, no thioether signal was observed in the *F*_obs_^interleaved-dark^ – *F*_obs_^laser-off^ [Figure 4(*g*)]. This lack of thioether bond signal confirms that there was no detectable light contamination. At the same time, the *F*_obs_^interleaved-light^ – *F*_obs_^laser-off^ [Figure 4(*h*)] and *F*_obs_^interleaved-light^ – *F*_obs_^interleaved-dark^ [Figure 4(*i*)] both show the expected electron density signature for the covalent adduct formation. Therefore, the *opaque* chips with the constrained (*flat* side) incidence of the laser light onto the sample did prove to be a successful setup in this experiment.

The reason for the success of the experiment in the *flat* orientation compared to the *open* was likely due to the reduced potential exposure of the crystals in this orientation (Figure 5). As stated in the methods, due to issues at the beginning of beamtime, the laser beam profile could not be precisely measured, only approximately inferred from scattered light off a black film. This means that the laser profile could conceivably be larger than the 50x50 μm^2^ estimate. Figure 5(*a*) shows how an increased laser profile could enable light contamination in the *open* chip orientation. The *flat* orientation could, by restricting the view of the crystals by the pump laser, help to prevent the contamination even with a larger laser profile. It is also possible that stray scattered light of either the collimator or chip sealing film may have been a contaminating factor [Figure 5(*b*)]. Another potential contributing factor was the synchronization of the stages and XFEL pulse. Subsequent experiments have showed that the XFEL pulse was delivered approximately 1 ms (12 μm) behind its intended location.

**Figure 5.**
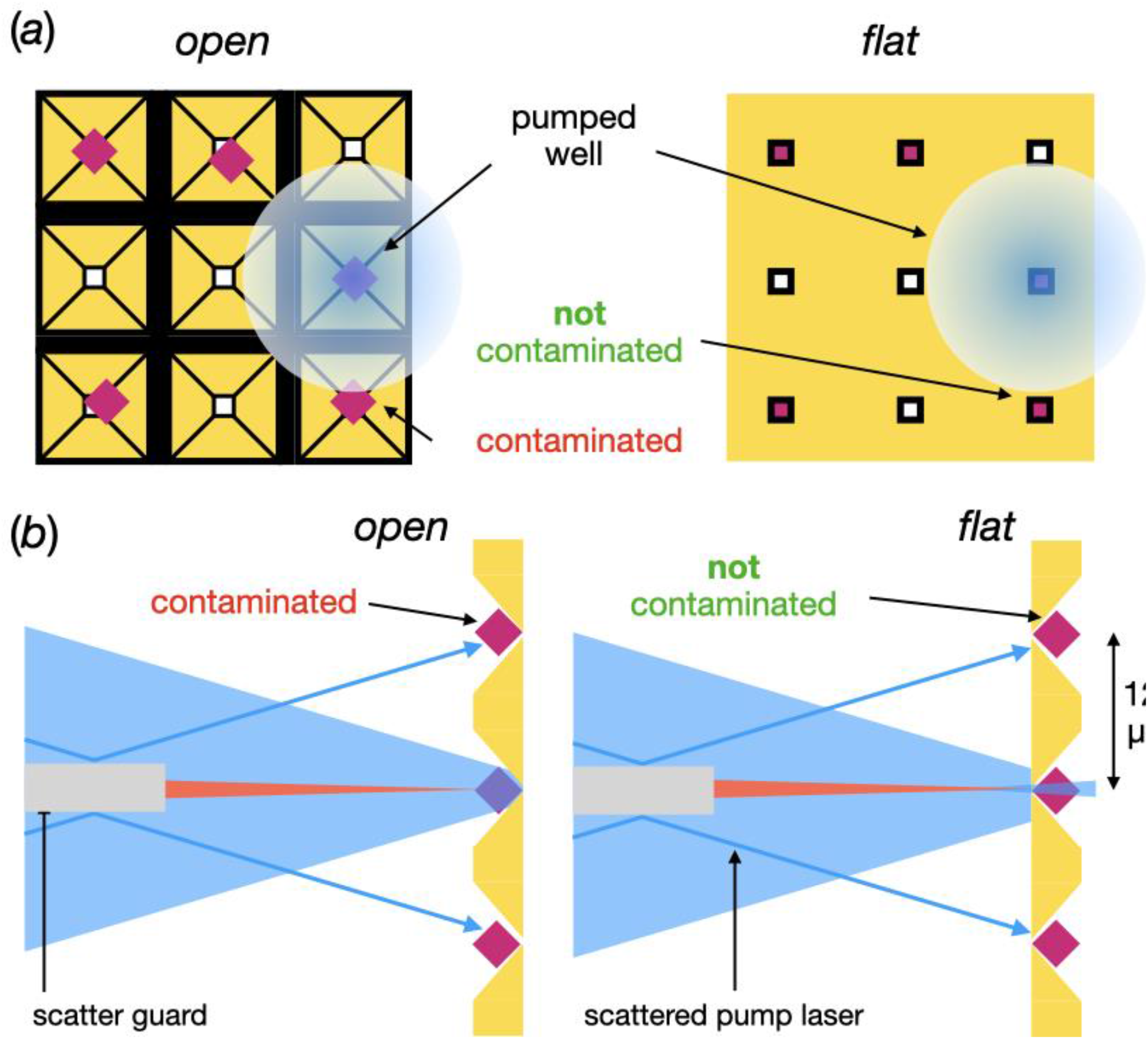
Possible explanations for the light contamination observed in the *open* orientation but not in the *flat*. (*a*) Schematic of the X-ray/pump laser view of the chip in the *open* and *flat* orientation. In the *open* view, the entire crystal is visible, enabling contamination *via* a large laser profile. The same large laser profile in the *flat* orientation does not give rise to contamination due to the restricted view of the crystals. (*b*) A schematic showing how potential scattered pump laser from the pre-sample scatter guard could give rise to light contamination. Again, restricted view of the crystals in the *flat* orientation prevents contamination of crystals in adjacent well.

### Reduction in sample consumption

Sample preparation is a laborious and challenging task for TR-SFX experiments. Various sample parameters must be considered and optimized depending on the delivery system used. Not only does the sample need to be of high quality but also the large quantities of crystalline protein can require months of sample production in preparation for every experiment. One of the attractive features of fixed-targets is their low sample consumption when compared to other delivery methods. The sample consumption from our fixed-target TR-SFX experiment was calculated to be only ∼200 μg for collection of every 10,000 indexed diffraction images. For comparison, Table 2 shows the quantities of samples consumed in several jet-based experiments. The development of the high-viscosity extrusion (HVE) for sample delivery was a significant improvement (>10x) when it comes to sample consumption compared to the first GDVN experiments. The HVE has resulted in a drastically increased number of successful TR-SFX experiments reported (Nogly *et al*., 2018, Mous *et al*., 2022, Nango *et al*., 2016, Suga *et al*., 2017, Nogly *et al*., 2016, Skopintsev *et al*., 2020, Claesson *et al*., 2020, Nass Kovacs *et al*., 2019, James *et al*., 2019, Hosaka *et al*., 2022, Liu *et al*., 2022, Maestre-Reyna *et al*., 2022, Li *et al*., 2021)

**Table 2.**
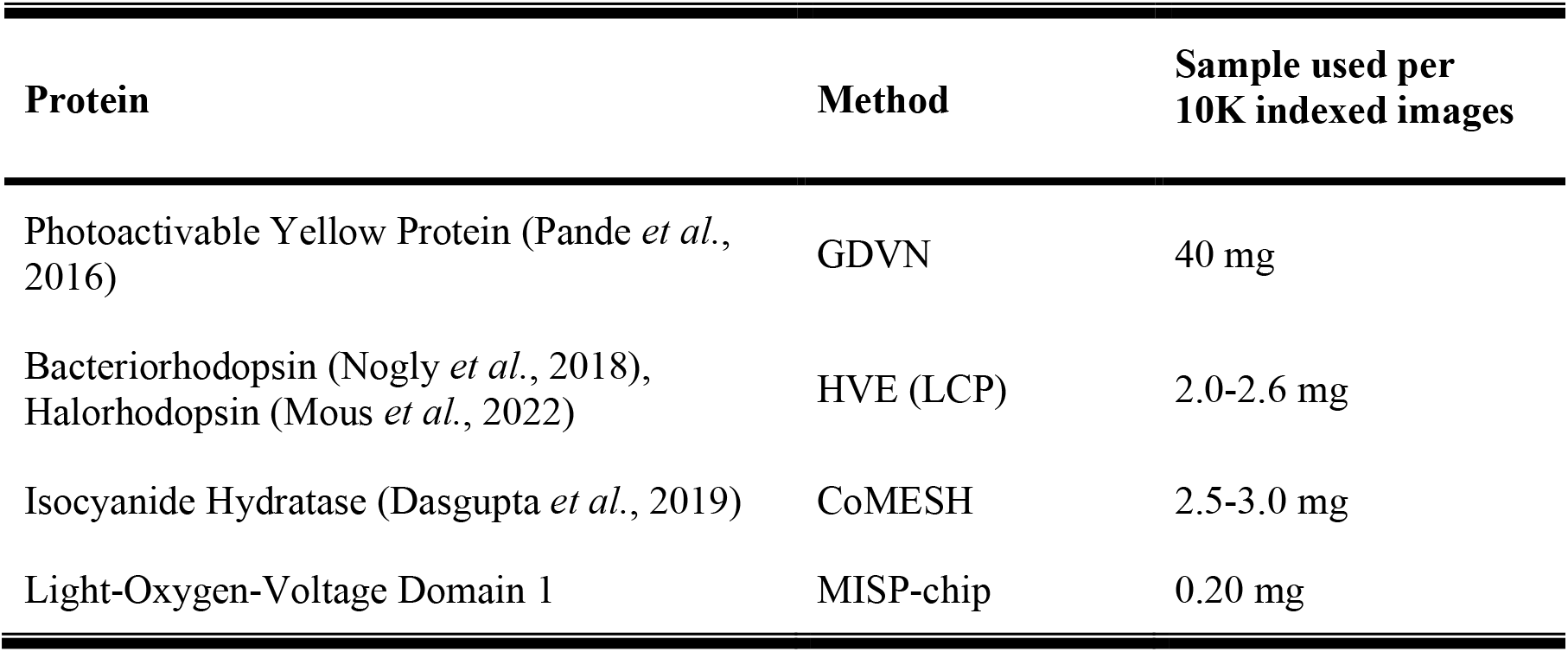
Pump-probed experiments sample consumption.

| **Protein** | **Method** | **Sample used per**<br>**10K indexed images** |
| --- | --- | --- |
| Photoactivable Yellow Protein (Pande *et al.*, 2016) | GDVN | 40 mg |
| Bacteriorhodopsin (Nogly *et al.*, 2018),<br>Halorhodopsin (Mous *et al.*, 2022) | HVE (LCP) | 2.0-2.6 mg |
| Isocyanide Hydratase (Dasgupta *et al.*, 2019) | CoMESH | 2.5-3.0 mg |
| Light-Oxygen-Voltage Domain 1 | MISP-chip | 0.20 mg |

The use of the MISP-chip reported here required approximately 10 times less sample than the HVE with less than 1 mg of protein required for a full dataset. With every improvement in sample delivery, TR-SFX becomes approachable to the broader scientific community and a wider range of interesting targets; many of which may not be easily overexpressed in greater quantities. Another practical point is that the preparation for the experiments can now be shortened in many cases from months to weeks facilitating timely completion of the TR-SFX projects.

The potential drawback of the sample delivery on the MISP-chips includes the time that it takes mounting each chip in a humid environment and transporting the chips in a humidity chamber from the dark room to the beamline, a procedure that took around half an hour for each 5 mounted chips. However, there is also a potential to employ robotic systems for the chip mounting for more efficient use of the beamtime. The light contamination is an important concern when compared to jet-based experiments, where the sample is enclosed in dark before delivery and injected in a stream. However, as shown in this commissioning experiment, even with a non-ideal chip configuration and large laser spot profile, a light contamination-free TR-SFX can still be performed.

The clear advantage of sample delivery fixed-target systems is lack of the often-tricky control of jet streams in the vacuum. Moreover, there isn’t a need for a dust-free environment when preparing the samples and crystal size distribution does not impact the efficiency of sample delivery. There are no additional sample losses related to clogging or instabilities (not accounted for in Table **2**) often observed with jet delivery.

All in all, we completed the TR-SFX experiment with limited sample usage but achieved the best diffraction limit for these crystals to date, making the approach a method of choice for many projects.

## Conclusion

XFELs have allowed for the rapid growth of time-resolved structural experiments which provide crucial information on the function of biological machines’ and molecular mechanisms. This is the first experiment of its type using a light-activated protein *Cr*LOV1 at the Cristallina endstation of the SwissFEL and using the novel MISP-chips for sample delivery. The light contamination present with the transparent MISP-chip was interesting to observe and showed the absolute necessity for using an opaque chip. With the opaque chips, the laser spot size and various other parameters are still critical to the success of the experiment. Many of these, particularly the laser spot profile, undoubtably lead to the contamination observed in the *open* orientation. Future commissioning experiments will focus on enabling TR-SFX in the *open* orientation as this conformation has the greatest propensity for crystal excitation. Ultimately, the reported experiments are an important step towards a making XFEL fixed-target sample delivery compatible with pump-probe time-resolved experiments and making these experiments more sample efficient. Increasing sample and experimental time efficiency will make these experiments more attainable for the general structural biology community.

## Acknowledgements

Project financed under Dioscuri, a program initiated by the Max Planck Society, jointly managed with the National Science Centre in Poland, and mutually funded by the Polish Ministry of Education and Science and the German Federal Ministry of Education and Research. This research was funded by National Science Centre, grant agreement No. UMO-2021/03/H/NZ1/00002 to P.N. We also acknowledge infrastructural support by Strategic Programme Excellence Initiative at Jagiellonian University - BioS PRA. For the purpose of Open Access, the author has applied a CC-BY public copyright licence to any Author Accepted Manuscript (AAM) version arising from this submission. Furthermore, this project was initiated with funding received from the Swiss National Science Fundation Ambizione grant PZ00P3_174169 to P.N. M.C. is supported by a studentship grant of the Swiss Nanoscience Institute, Project #1904. This project has also received funding from the European Union’s HORIZON 2020 Research and Innovation Program under the Marie S. Curie grant agreement No. 701647. We acknowledge the Paul Scherrer Institute, Villigen, Switzerland for provision of free-electron laser beamtime at the Cristallina experimental station on the SwissFEL ARAMIS branch. Cristallina was realized with financial support from the Swiss National Science Foundation and the University of Zürich under Project Nr. 206021_183330.

